# Revealing the cell-material interface with nanometer resolution by FIB-SEM

**DOI:** 10.1101/123794

**Authors:** Francesca Santoro, Wenting Zhao, Lydia-Marie Joubert, Liting Duan, Jan Schnitker, Yoeri van de Burgt, Hsin-Ya Lou, Bofei Liu, Alberto Salleo, Lifeng Cui, Yi Cui, Bianxiao Cui

## Abstract

The interface between biological cells and non-biological surfaces profoundly influences cellular activities, chronic tissue responses, and ultimately the success of medical implants. Materials in contact with cells can be plastics, metal, ceramics or other synthetic materials, and their surfaces vary widely in chemical compositions, stiffness, topography and levels of roughness. To understand the molecular mechanism of how cells and tissues respond to different materials, it is of critical importance to directly visualize the cell-material interface at the relevant length scale of nanometers. Conventional ultrastructural analysis by transmission electron microscopy (TEM) often requires substrate removal before microtome sectioning, which is not only challenging for most substrates but also can cause structural distortions of the interface. Here, we present a new method for *in situ* examination of the cell-to-material interface at any desired cellular location, based on focused-ion beam milling and scanning electron microscopy imaging (FIB-SEM). This method involves a thin-layer plastification procedure that preserves adherent cells as well as enhances the contrast of biological specimen. We demonstrate that this unique procedure allows the visualization of cell-to-material interface and intracellular structures with 10nm resolution, compatible with a variety of materials and surface topographies, and capable of volume and multi-directional imaging. We expect that this method will be very useful for studies of cell-to-material interactions and also suitable for *in vivo* studies such as examining osteoblast adhesion and new bone formation in response to titanium implants.

---

Many biological applications and biomedical devices require direct contact between biological cells and non-biological materials^1^. In the case of medical implants, the cell-to-material interface is a key determinant for successful device integration with surrounding tissues, providing mechanical support and minimizing host foreign body responses^2^. In addition to providing mechanical support, non-biological materials are actively explored for inducing the regeneration and repair of surrounding tissues^2^. In this context, the cell-to-material interface is essential in regulating cell signaling, guiding cell migration, and controlling stem cell differentiation and lineage specificity^3,4^.

To date, ultrastructural analysis by transmission electron microscopy (TEM) is the most detailed method for analyzing the cell-to-material interface. TEM can resolve the cell membrane and subcellular structures, which reveals how cells make contacts with the substrate surface and provides accurate measurement of the gap distance between the cell membrane and the material surface^5–7^. However, the TEM method requires embedding the sample in millimeter-sized resin blocks and, then, sectioning them into ultra-thin slices (∼100 nm thickness) with mechanical knives. In many cases (*i.e*. hard materials such has glass and metals), the substrate has to be removed by chemical etching or physical separation before sectioning. Substrate removal is often not feasible for many systems, and even if feasible, the procedure can induce structural artifacts at the interface. Furthermore, in TEM resin-blocks, the context of the cell is lost unless a 3D reconstruction is carried out after a tedious procedure of sectioning, sorting and imaging hundreds of individual slices.

A combination of focused ion beam (FIB) and scanning electron microscopy (SEM) constitutes an alternative approach for sectioning/imaging of materials^8^ and biological specimens^9^. Unlike TEM, FIB-SEM allows *in situ* ion-based milling of the specimen to reveal interfaces at any desired location. The use of FIB-SEM for examining the interaction of cells with engineered surfaces has been previously explored by us^10,11^ and by others^12–17^. However, using FIB-SEM for cell-to-material interface studies is severely limited by structural damages and the poor contrast of the biological specimen, usually prepared by hard drying procedures. The drying procedure can induce substantial volume shrinkage^18,19^, as well as cavities (sponge-like morphology) in place of the intracellular compartments^14,16^. The lack of contrast in biological specimens results in an inability to resolve the intracellular structures or the cell membrane. Recently, a thin-layer resin embedding method has been developed ^16,20^, but the lack of contrast (*i.e*. no heavy metals) does not allow the visualization of the plasma membrane or intracellular compartments at the interface. In this work, we present a new FIB-SEM method that is capable of *in situ* visualization of the cell-to-substrate interface with high contrast that resolves subcellular structures and the cell-to-material junction with 10 nm resolution. The method is compatible with diverse materials (quartz, doped silicon, conductive polymer) and various surface topographies, allowing clear identification of how the cell membrane interacts with nanoscale features (protrusions and cracks) on the substrate.

At the core of our FIB-SEM method, there is a new sample preparation method based on controlled thin-resin plastification of adherent cells with heavy metal staining and as well as preservation of the contacting material. Unlike the usual hard drying method, this method embeds cells in a thin plastic layer, which not only preserves the subcellular structures but also provides a solid support for the subsequent FIB milling. The thin-layer plastification method includes five major steps: cell fixation, heavy metal staining, resin infiltration, extracellular resin removal, and resin polymerization (Figure 1a). First, mammalian cells cultured on the desired substrate (doped silicon, polymer or nanostructured quartz) are fixed by glutaraldehyde to crosslink intracellular structures (*i.e*. proteins) so that they can withstand the subsequent staining and embedding processes. After fixation, the cells are treated with an osmium series (RO-T-O procedure^21,22^) and *en bloc* staining (see **Methods**), which not only provide high contrast to membrane and protein structures, but also help to preserve lipids in subsequent steps. Then, the cells are infiltrated with liquid epoxy-based resin. Traditional resin embedding procedures for TEM typically result in a 2 - 5 millimeter-thick polymer block, preventing the visualization of the whole-cell morphology. In our method, after resin infiltration and before resin polymerization, a resin-removal step is introduced that strips off excess extracellular resin by draining and flushing the sample with ethanol. This step thins down the resin coating outside the cell membrane to tens of nanometers while maintaining a stable intracellular resin embedding^20^. The final step involves curing the liquid resin to a thin layer of plastic with cells embedded inside. Since extracellular resin is largely removed, cell topography and membrane protrusions in contact with the underlying substrate are clearly visible under SEM. Figure 1b shows a HL-1 cell cultured on a quartz substrate with arrays of quartz nanopillars, and **Supplementary Figure S1** shows PC12 cells and primary cortical neurons cultured on flat glass substrates, where fine features on the cell membrane are well preserved.

**Figure 1:**
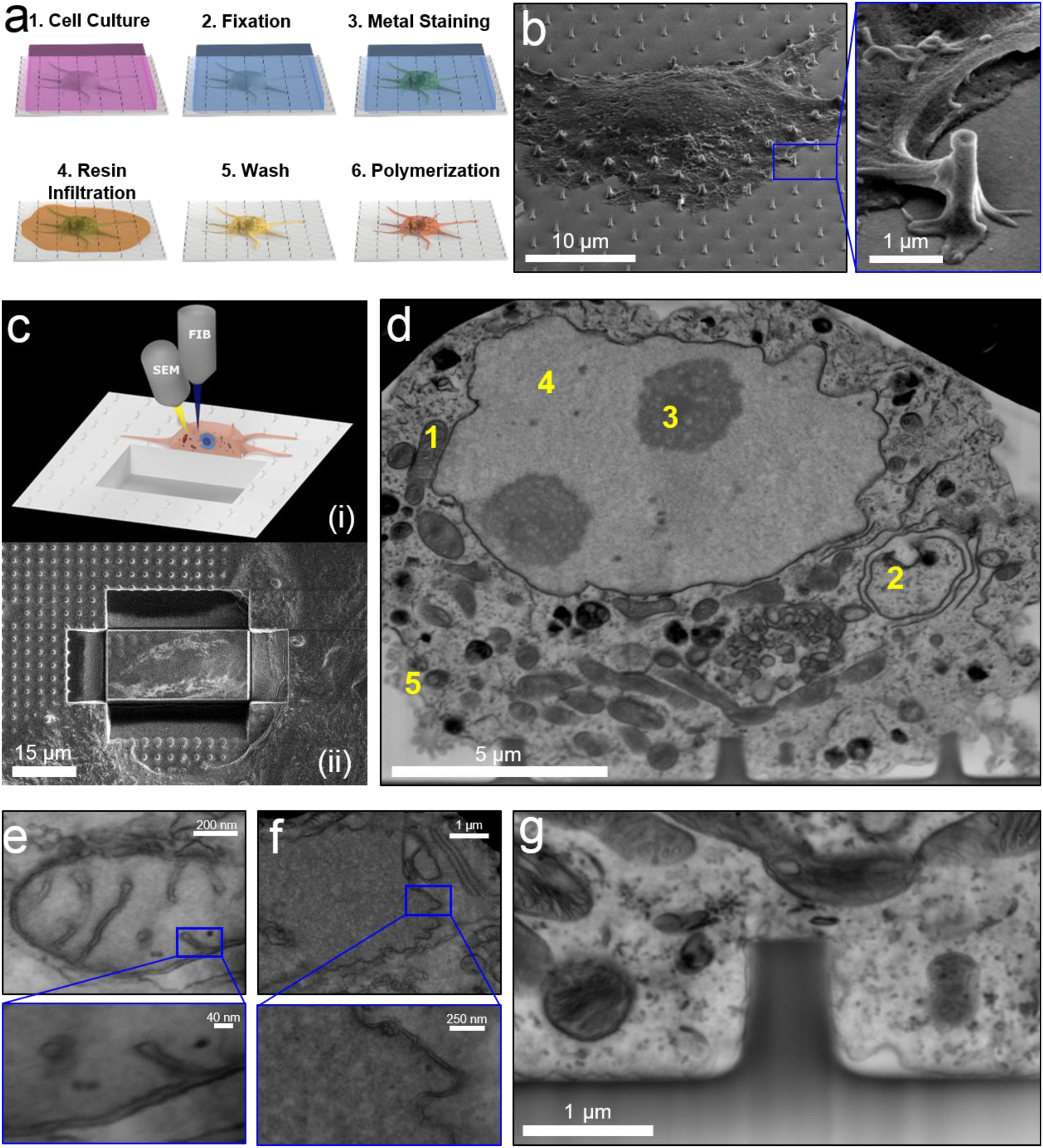
Imaging the cell-to-material interface by FIB/SEM. **a)** Schematics of the sample preparation procedure by thin-layer resin plastification with contrast enhancement. **b)** A SEM image of a plastified HL-1 cell on a quartz substrate with nanopillars clearly shows that extracellular resin is removed and the cell morphology is clearly visible. The insert shows that the membrane protrusions in contract with a nanopillar are well preserved. **c)** Schematics (i) and experimental results (ii) of using FIB milling to cut trenches through the cell and the substrate and open up the interface. **d)** A SEM image of the interface after FIB milling reveals intracellular compartments and organelles such as mitochondria (1), intracellular membranes (2), nucleoli (3), nucleus (4) and cellular membrane (5). **e-f)** Zoomed-in FIB-SEM images of mitochondria (e) and nuclear envelope (f), The insets clearly resolves the inner and outter membranes and interstitial space. **g)** At the interface between the cell and the quartz substrate, the plasma membrane is shown to warp around a vertical nanopillar. Intracellular structures and local curvatures on the plasma membrane resembling clathrin-mediated endocytosis events can be clearly identified. Figures **d-g** have been acquired from backscattered detectors (voltage:5-10 kV, current: 0.64-1.4 nA), tilt is 52°.

Samples prepared *via* thin layer-plastification are directly mounted on FIB-SEM for *in situ* examination of the cell-to-substrate interface. For this purpose, we first examine a large sample area by SEM to identify locations of interest, such as places where cell membranes are in contact with the surface features like nanopillars. Once a desired area is located, it is coated with a thin layer of platinum to facilitate the dissipation of ions and prevent sample damage during the next FIB milling step (see **Methods** and **Supplementary Figure S2**). Then, a high-energy gallium ion beam is focused on the sample to cut through the platinum protection layer, the cell-embedded thin plastic layer underneath, and at least 1 μm deep into the substrate. This process is repeated to remove material and open up a vertical surface (Figure 1c). Then, a low-current, e.g. 80 pA, ion beam is used to remove re-deposited material and polish the cross section. This step is critical for limiting the curtaining phenomena and ion-induced structural damage at the interface^14^. SEM visualization of the cross section shows intracellular space and the interface between the cell membrane and the substrate. Unlike previous FIB-SEM images that usually contain sponge-like structures with no discernable subcellular structures^15,16,20^, our FIB-SEM image shows very clear subcellular structures such as the cell membrane, the nucleus, nucleoli, the nuclear envelope, mitochondria, lysosomes, and multi-vesicular bodies (Figure 1d). To be consistent with published TEM images, all FIB-SEM images are black-and-white inverted. Original images are shown in **Supplementary Figure S2**.

To determine the resolution capabilities of our FIB-SEM method, we have examined a group of well-characterized cellular compartments using high magnification SEM imaging. Figure 1e shows an image of a mitochondrion that clearly resolves inner and outer membranes (∼10 nm distance) as well as the cristae structures defined by the inner membrane. Figure 1f shows the structure of nuclear envelope with clearly-resolved inner and outer membranes with an interstitial space of about 20 nm. Endoplasmic reticulum (ER) structures as parallel running membranes can be seen in the vicinity of the nucleus, and the associated small granules attached to the membrane of the ER likely are ribosomes (**Supplementary Figure S3a**). Other intracellular structures have been resolved such as multi-vesicular bodies and endocytic vesicles (**Supplementary Figure S3b-d**). Furthermore, FIB-SEM clearly reveals that the plasma membrane is very close (< 50 nm) to the flat substrate surface and contours around local nanopillar features (Figure 1g).

Our FIB-SEM method is compatible with substrates with different surface topographies and different materials, *i.e*. a quartz substrate with nanotubes (**Supplementary Figure S4**). As shown in Figures 2a, i&ii, the cell membrane attaches tightly to the flat areas of the quartz substrate and wraps around the outside surface of a nanotube, with intracellular structures in the vicinity clearly visible. However, inside the nanotube, the cell membrane did not conformably follow the surface contour and only extended into the top part of the hollow center (Figure 2a, iii) as previously observed by TEM^23^. In addition to the quartz substrate, we have demonstrated that our FIB-SEM method can be used for substrates made with the conductive polymer blend Poly(3,4-ethylenedioxythiophene):Polysterene Sulfonate (PEDOT:PSS) and doped-silicon (conductive), with all surface were coated with poly-L-lysine to facilitate cell culture. The surface of the PEDOT:PSS is patterned into parallel grooves (Figure 2b, i & **Supplementary Figure S4**). The cell appears well spread on the PEDOT surface, however the FIB-SEM image reveals that the cell membrane is much further away from the PEDOT substrate (∼ 100 nm average on flat areas) than from the quartz substrate (∼25 nm average on flat areas). Locations where the cell membrane made close contacts with the PEDOT substrate can be clearly identified (arrow heads in Figure 2b, ii). The cell membrane extends into the patterned groove (Figure 2b, ii) and into some local cracks on the substrate (Figure 2b, iii). The surface of the doped-silicon substrate has randomly distributed nanocone features (Figure 2c, i and **Supplementary Figure S4**). By FIB-SEM, we observed that the cell membrane is far from the flat substrate (∼200 nm) in most places, but makes close contacts with the top of the nanocones.

**Figure 2:**
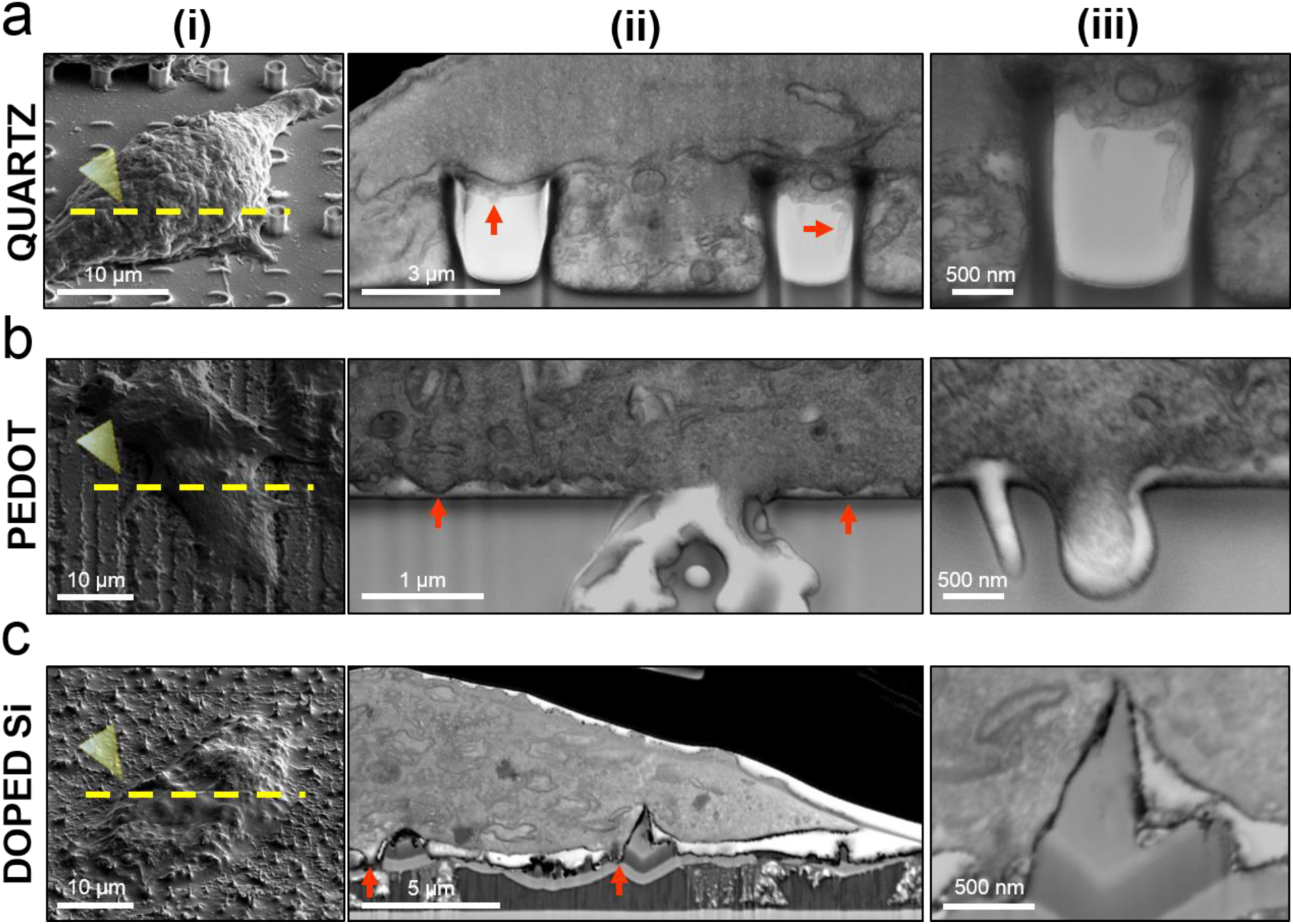
FIB/SEM imaging of the interface is compatible with a variety of materials and surface topography. **a)** SEM images of cells cultured on a quartz substrate with nanotubes, before (i) and after FIB milling (ii, iii). The interface between the cell and the quartz nanotube shows that the plasma membrane not only warps around the outsize of the nanotube and but also extends into the top part of the nanotube cavity (red arrows and iii). **b)** SEM images of cells cultured on grooved PEDOT:PSS surface before (i) and after FIB milling (ii); The cell membrane is further away from the PEDOT surface than the quartz surface. Red arrows in (ii) indicate attachment points of the plasma membrane. Image in (iii) shows the membrane protruding into a pit on substrate. **c)** SEM images of cells on a doped-silicon substrate with randomly distributed nanocones before (i) and after FIB milling (ii). Zoom-in image (iii) shows the plasma membrane partially wraps around the nanocone walls through attachment points (red arrows in ii). Figures (i) have been acquired by a secondary electron detector, (ii) and (iii) have been acquired with a backscattered detector (voltage: 5 - 10 kV, current: 0.64 - 1.4 nA). Tilt is 52° in all images.

Since FIB-SEM allows repetitive milling and imaging, it is possible to image a volume of interest (VOI) at high resolution (Figure 3a). We used low current (e.g. 80 pA) for sequential FIB milling, which achieves slice thickness of about 39 nm and well beyond the capability of mechanical slicing by means of ultramicrotomes (70 - 200 nm). Figures 3b&c show two representative cross-sections of the same cell (shown in Figure 3a) interacting with two different lines of nanopillars. By sequentially imaging a set of 72 sequential sections, we reconstructed a 3D intracellular space and its interaction with nanopillars using a segmented 3D reconstruction method (Figure 3d, **Supplementary Movie 1**). In particular, we modeled the 3D morphology of the nuclear envelope, nucleoli, and the non-adherent cellular membrane domain, which were individually constructed and overlaid to the remaining structures as shown in Figure 3e. The nuclear envelope appears to be bend upward on top of a nanopillar for as much as 800 nm (Figure 3f), agreeing well with our previous observation by TEM^24^.

**Figure 3.**
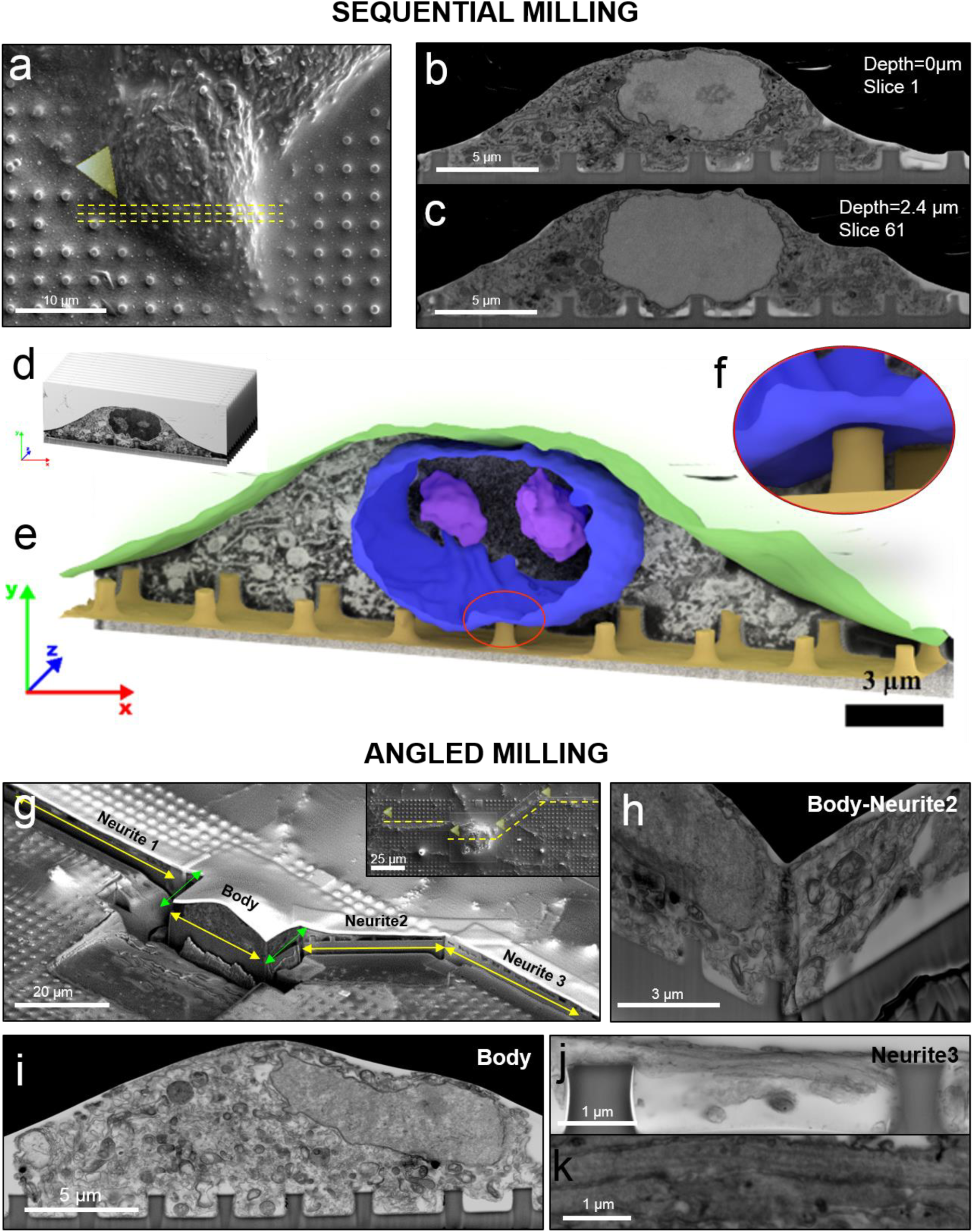
FIB-SEM for sequential volumetric imaging and multi-angled imaging. **a)** A SEM image of a plastified HL-1 on nanopillars where yellow dashed lines indicate the region of interest for the sequential milling. **b-c)** SEM images of two exemplary slices from a stack of 78 slices at two different pillars’ lines. d) Images collected in the stack were assembled, segmented, and analyzed. **e)** Automated 3D reconstruction of membrane and nuclear envelope were overlaid to SEM background image. f) Reconstruction shows that the nuclear envelope is deformed upward by a nanopillar. **g)** FIB milling of a neuron where yellow arrows indicates the regions of interest and green lines indicates the connecting regions (the inset shows a SEM image of the same neuron before FIB milling). **h)** A FIB-SEM image of the body-neurite 2 connecting region opened at 90-degree angle. **i)** A FIB-SEM image of the neuronal body on a line of nanopillars. **j)** A FIB-SEM image of neurite 3 on top of nanopillars. **k)** Zoomed-in image of neurites reveals multiple longitudinally orientate microtubules parallel to the direction of the neurite.

Unlike the ultramicrotome sectioning method that slices materials sequentially in only one direction, the FIB-SEM method is highly versatile and allows sectioning of the same sample with different directions at multiple locations. This capability is often important for cells with protrusions such as neurons. Primary cortical neurons from embryonic rats were cultured on a quartz substrate with arrays of solid nanopillars. After 5 days of culturing *in vitro*, neurons were fixed and processed for FIB-SEM imaging as described earlier. A SEM image in Figure 3g **(insert)** shows a neuron cell body together with multiple neurites growing out from the cell body. We first identified four regions of interest from the SEM image: the cell body, neurite-1, neurite-2 and neurite-3. Then, after coating a layer of Pt, FIB milling was used to cut open the interfaces along six connecting lines (yellow arrowed lines corresponding to four regions of interest and green arrowed lines being the connecting lines in Figure 3g). FIB-SEM imaging of the cell body shows the nuclear, large number of intracellular organelles and the plasma membrane wrapping around nanopillars (Figure 3i). By multiple angle milling, FIB-SEM also offers a unique advantage of examining a location from multiple directions as shown by the 90-degree intersection between the neurite-2 and the cell body (Figure 3h). The cross-section of neurite-3 is shown in Figure 2j, which illustrates a neurite attached to the top and the side of two nanopillars. A magnified image of a neurite reveals multiple longitudinally orientated microtubules parallel to the direction of the neurite (Figure 3k), comparable in morphology to those investigated by TEM^25,26^. FIB-SEM images of Neurite-1 and Neurite-2 connected to the cell body are shown in **Supplementary Figure S5**.

Furthermore, we show that our FIB-SEM method is suitable for correlating light and electron microscopy images (CLEM). For this purpose, we first proved that for cells fixed and stained with fluorescent phalloidin for actin filaments and immunostained for clathrin (**Supplementary Figure S6**). The fluorescence image taken before the resin infiltration step shows actin accumulation on nanopillars (**Supplementary Figure S6**), agreeing with results previously reported^24^. Using nanostructures as location markers, the subsequent resin embedding and SEM imaging shows the cell morphology perfectly correlating with the fluorescence imaging, which further confirms that the cell volume was well preserved without any significant alteration.

Finally, we demonstrate that our FIB-SEM method is also compatible with correlative microscopy of living cells transfected with a APEX2-GFP-based construct. Recently, APEX2, a genetically encoded peroxidase, has been used to selectively enhance the contrast for certain subcellular structures under TEM^27,28^ (e.g. mitochondria). We constructed an APEX2-GFP-CAAX fusion plasmid that selectively targets APEX2 to the plasma membrane to further enhance the contrast at the cell-to-material interface. We transfected cells growing on nanopillars with APEX2-GFP-CAAX allowing initial localization of transfected cells by live fluorescence imaging (Figure 4a). Then, after cell fixation and before osmium staining, 3,3-diaminobenzidine (DAB) and H_2_O_2_ are added to the solution, where APEX2 catalyzes the polymerization and deposition of DAB in its vicinity. The polyDAB recruits osmium in the subsequent staining step to give greater contrast to APEX2-labeled structures. After thin-resin plastification, the cell can be visualized by SEM (Figure 4b) or ion microscopy (**Supplementary S7**). The FIB milling at the location of interest opens the cross section for examining the cell-to-substrate interfaces (Figure 4c). We compared the interfaces for APEX2-GFP-CAAX transfected cells *vs*. non-transfected cells in the same culture. As shown in Figures 4d&e, under identical conditions, APEX2-CAAX transfected cells show a higher membrane contrast than non-transfected cells (the plasma membrane is visible in both cases). We note that the APEX2-based contrast enhancement is less distinct than expected^27^, likely because we used uranyl acetate staining that is known to have a higher affinity to osmium in membranes than to DAB polymers. Nevertheless, our method uniquely allows direct correlation between living cells in fluorescence, SEM after thin plastification and the cross section after selectively FIB milling.

**Figure 4.**
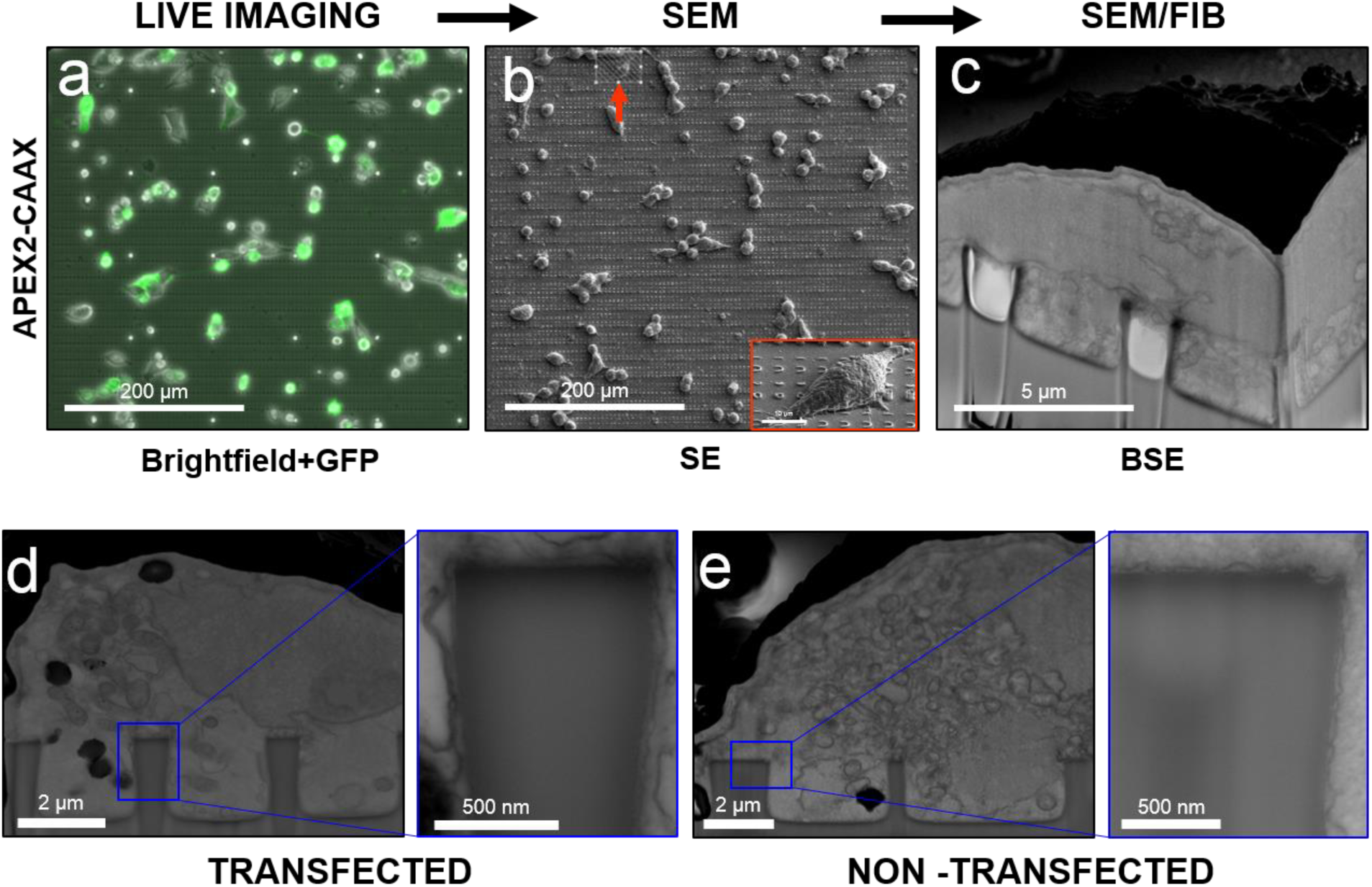
Enhancing FIB/SEM imaging by a APEX2 tag. **a)** Overlay of the brightfield and the fluorescence (GFP) images of cells transfected with APEX2-GFP-CAAX. **b)** A SEM image of the same area after thin-layer plastification. The arrow shows the target cell before FIB milling (52° tilted SEM image in the inset). **c)** The FIB-SEM image of the target cell opened at 90 degree. **d)** A FIB-SEM image of a APEX2-GFP-CAAX transfected cell. The inset shows the membrane contrast at the interface. **e)** A FIB-SEM image of a non-transfected cell in the same culture, imaged at the same condition. The inset shows the plasma membrane with lower contrast around the nanopillar.

In conclusion, we demonstrated a new FIB-SEM method for imaging of the cell-to-material interface *in situ*, without removing the substrate. This method achieves a contrast and resolution higher than TEM previously used for similar investigations, Moreover, for the first time cells interfacing materials such as PEDOT:PSS have been investigated besides more common materials such as quartz and silicon-based surfaces with various topology. Our FIB-SEM method has unique advantages of examining a large sample area of an artefact-free plastified cell, opening up cross sections at any desired location, achieving volume reconstruction, performing multi-directional milling, and compatible with correlative microscopy and APEX2-based enhancement. In perspective, our method can be used for any cell-material interaction investigation so that, for example, the interaction of cells/tissue with medical devices.

## Acknowledgments

The authors thank the Stanford NanoShared Facility (SNSF) for a seed grant to a complimentary use of the Helios i600 and Dr. Juliet Jamtgaard and Dr. Richard Chin for the useful discussions. The authors also acknowledge the National Science Foundation for the grants NSF 1055112 and NSF 1344302.

## Methods

### • Nanostructures fabrication, characterization and preparation

#### Fabrication and characterization of quartz nanopillars

Nanostructures (NSs) used in this work were fabricated on 4-inch quartz wafer using electron-beam lithography (EBL). In brief, the wafer was diced into pieces of 2 cm × 2 cm square. After sonication cleaning in acetone and isopropanol, the pieces were spin-coated with 300 nm of ZEP-520 (ZEON Chemicals), followed by E-Spacer 300Z (Showa Denko). Desired patterns were exposed by EBL (Raith150) and developed in xylene. The mask was then created by sputter deposition of 100 nm Cr and lift-off in acetone. NSs was generated by reactive ion etch with CHF_3_ an O_2_ chemistry (AMT 8100 etcher, Applied Materials). Before cell culture, the substrate was cleaned in O_2_ plasma and immersed in Chromium Etchant 1020 (Transene) to remove Cr masks. SEM (FEI Nova) imaging was performed on 3 nm Cr sputtered substrates to measure the dimension of different NSs.

#### Silicon-nanocones

A monolayer polystyrene nanosphere (PS) array, which consists of PSs with an averaged diameter of 3 μm, were self-assembled on glass-based silicon substrates with a Langmuir–Blodgett method. To control the effective intervals between the formed silicon nanopillars, a reactive ion etching (RIE) process with oxygen (O_2_) as an etching gas was then followed to shrink PSs (with a final diameter of 1 μm). Silicon nanopillars were lastly formed on glass substrates by introducing chlorine (Cl_2_) and hydrogen bromide (HBr) gasses to reactive-ion-etch the silicon materials exposed to the plasma.

#### PEDOT:PSS nanogrooves

Fused silica glass substrates were cleaned using a standard soap, acetone, isopropanol sonication sequence. Poly(3,4-ethylenedioxythiophene) polystyrene sulfonate (PEDOT:PSS) (Heraeus, Clevios PH 1000) solution in water was doped with 5 wt % ethylene glycole, 0.1 wt% Dodecyl Benzene Sulfonic Acid (DBSA) as a surfactant and 1 wt% (3-glycidyloxypropyl)trimethoxysilane (GOPTS) as a crosslinking agent to improve film stability. EG, DBSA and GOPS were all obtained from Sigma Aldrich. After spin-coating at 1000 rpm for 2 mins the films where baked at 120 °C for 10 mins. The nanogrooves were created by focusing a Ti:Sapphire femtosecond laser (Spectraphysics, 100 fs, 50 mW) to a 5 μm spot right above the surface of the PEDOT:PSS film and scanning the beam over the films at 2 mm/s.

#### Sample preparation for cell culture

Quartz substrates were treated with Pirana solution with sulfuric acid and hydrogen peroxide (Fisher Scientific), in a 7:1 dilution at room temperature overnight. Samples were washed with distilled water, dried and placed in 70% ethanol in a sterile hood. Samples were washed with sterile distilled water and allowed to dry. After a 15 mins UV light exposure, samples were incubated overnight with 0.01% poly-L-lysine (Sigma Life Science) for primary neurons and HEK cells cultures or with 1 mg/ml fibronectin (Life Technologies) in 0.02% gelatine solution for HL-1 cells. COS-7 cells were directly plated on the substrate after sterilization.

### • Cell culture and transfection

#### Primary neurons

Cortices were extracted from rat embryos at embryonic day 18 and incubated with 0.25% Trypsin/EDTA (Corning) in 33 mm Petri dish for 5 min at 37°C. The tissue-trypsin/EDTA solution was transferred into a 2 mL plastic tube. The tissue settled at the bottom of the tube and left over trypsin/EDTA was removed. Neurobasal^®^ media (Gibco) was supplemented with 1% B27 (Gibco), 0.25% glutaMAX, (Gibco) and 0.1% gentamycin antibiotic (Gibco). One ml of warm media was added and, then, the tube was gently swirled by hand. This procedure was repeated 5 times and after the last media exchange, the tissue was dissociated until resulting in a cell solution. 80,000 cells were suspended in 3 mL and placed on each substrate. The media was replaced completely 2 hrs after seeding time. Every second day, half of the media was exchanged with freshly-prepared warm (supplemented) Neurobasal^®^ media.

#### HL-1 cells

Confluent HL-1 cells, cultured in a 33 mm Petri dish, were incubated with 1 mL of 0.25% trypsin/EDTA for 5 mins at 37° C. The cell-trypsin solution was transferred into a 15 mL tube and 2 mL of Claycomb media (Sigma Life Science) supplemented with 10% of fetal bovine serum (Sigma-Aldrich), 100 μg/mL penicillin/streptomycin (Sigma Life Science), 0.1 mM norepinephrine (Sigma-Aldrich) and 2 mM glutaMAX were add. The cell solution was placed in a centrifuge for 3 mins with a rotation of 1300 rpm. Cells’ pellet was resuspended in 1 mL of media and 50 μl of the resuspension was plated on each substrate in addition to 3 mL of supplemented media.

#### HEK 293 cells

HEK 293 expressing channels Na_V_ 1.3 and K_IR_ 2.1 were acquired by Adam Cohen laboratory and maintained in DMEM/F12 (Gibco), 10% FBS (Gibco), 1% penicillin/streptomycin (100 μg/mL, Gibco), geneticin (500 μg/mL, Gibco) and puromycin (2 μg/mL, Fisher Scientific). At 80% confluency, cells were divided, re-suspended and plated on quartz substrates as for HL-1 cells.

#### COS-7 and U2OS cells

Cells were maintained in DMEM supplemented with 10% fetal bovine serum and at 90% confluency they were divided as for HL-1 cells and plated on the substrates.

### • Ultra-thin plastification and RO-T-O procedure

Substrates with cells were rinsed with 0.1 M sodium cacodylate buffer (Electron Microscopy Sciences) and fixed with 3.2% glutaraldehyde (Sigma-Aldrich) at 4°C overnight. Specimens were then washed (3 × 5 mins) with chilled buffer and quenched with chilled 20 mM glycine solution (20 mins). After rinsing (3×5 mins) with chilled buffer specimens were post-fixed with equal volumes of 4% osmium tetroxide and 2% potassium ferrocyanide (Electron Microscopy Sciences, *RO* step) (1 hr on ice). Samples were then washed with chilled buffer (3× 5 mins) and the solution replaced with freshly prepared 1% thiocarbohydrazide (Electron Microscopy Sciences, *T* step) (20 mins at room temperature). After rinsing with buffer (2 × 5 mins), the samples were incubated with 2% aqueous osmium tetroxide (*O* step) (30 mins at room temperature. Cells were again rinsed (2 × 5 mins) with distilled water and then, finally, incubated with syringe-filtered 4% aqueous uranyl acetate (Electron Microscopy Sciences, *en bloc* step) (overnight 4°C). Cells were rinsed (3 × 5 mins) with chilled distilled water, followed by gradual dehydration in an increasing ethanol series (10%-30%-50%-70%-90%-100%, 5-10 mins each on ice). The last exchange with 100% ethanol solution was performed at room temperature. Epoxy-based resin solution was prepared as previously described(*19*), and samples infiltrated with increasing concentrations of resin in 100%ethanol, using these ratios: 1:3 (3 hrs), 1:2 (3 hrs),1:1 (overnight), 2:1 (3 hrs), 3:1 (3 hrs). Infiltration was carried out at room temperature and in a sealed container to prevent evaporation of ethanol. Samples were then infiltrated with 100% resin overnight at room temperature. The excess resin removal was carried out by first draining away most of the resin by mounting the sample vertically for one hour and, then, rapidly rinsing with 100% ethanol prior to polymerization at 60°C overnight.

#### APEX2 contrast enhancement

##### Plasmid preparation

To make APEX-GFP-CAAX, CIBN-GFP-CAAX (a gift from Dr. Chandra Tucker in University of Colorado Denver) was first linearized by NheI and AgeI restriction enzymes to remove CIBN. APEX fragment was amplified from Connexin-GFP-APEX (obtained from Addgene) with forward primer CGTCAGATCCGCTAGCGCCACCATGGGAAAGTCTTACCCAACTG and reverse primer CATGGTGGCGACCGGTACATGGGCATCAGCAAACCCAAGC. Using InFusion cloning kit (Clontech, Mountain View, CA, USA), APEX was inserted into the previously linearized backbone.

##### Transfection

Cells were transfected with APEX2-GFP-CAAX plasmids (340 ng) using Lipofectamine 2000 (Life Technologies) according to the manufacturer’s protocol. The transfected cells were allowed to recover and express the desired protein for 18 hrs prior to live imaging performed with a microscope LEICA DMI 6000B (Leica).

##### Osmication and staining

Cells were washed with 0.1 M sodium cacodylate buffer and fixed with 2.5% glutaraldehyde for at least 1 hr at 4°C. Substrates were washed three times with chilled buffer and quenched with chilled 20 mM glycine in buffer solution for 20 mins. Afterwards, cells were washed with chilled buffer followed by 3,3’-diaminobenzidine (Sigma-Aldrich) solution which had been freshly prepared as follows: DAB powder was mixed with 1 M HCl to reach a final concentration of 1.4 mM. Thereafter the DAB solution was mixed (equal volumes) to 0.03% H_2_O_2_ (in 0.1 M cacodylate buffer). Cells were osmicated with 2% osmium tetroxide for 1 hr at 4°C, washed with chilled buffer and incubated with 2% potassium ferrocyanide overnight at 4° for the reduced osmium procedure. For membrane enhancement in FIB-SEM, cells were treated with RO-T-O enhancement, ethanol dehydration and ultra-thin plastification as described above.

### • Scanning electron microscopy imaging and focused ion beam sectioning Sample preparation

Each sample was glued with colloidal silver paste (Ted Pella Inc.) to a standard stub 18 mm pin mount (Ted Pella Inc.). A very thin layer of gold-palladium alloy was sputtered on the sample before imaging.

#### SEM imaging

Samples were loaded into the vacuum chamber of a dual-beam Helios Nanolab600i FIB-SEM (FEI). For selecting a region of interest, an (electron) beam with accelerating voltage 3-5 kV, and current 21 pA - 1.4 nA, was applied. For image acquisition of whole cells (*i.e*. Figure 1b) a secondary electrons detector was used. For cross section imaging, a beam acceleration voltage of 2 kV - 10 kV was selected, with the current ranging between 0.17 - 1.4 nA, while using a backscattered electrons detector (immersion mode, dynamic focus disabled in cross section, stage bias zero), a dwell time of 100 μs and 3072 × 2048 pixel store resolution. For the sequential sectioning, the function iSPI was enabled in order to slice and acquire an image of the stack every 38.5 nm with 5 kV voltage, 1.4 nA current and 1024×884 resolution.

#### FIB sectioning

Regions of interest were preserved by electron-assisted deposition of a 0.5 μm double platinum layer, and ion-assisted deposition of a (nominal) 1μm thick coating. First, trenches were created with an etching procedure fixing an acceleration voltage of 30 kV and currents in the range 9.1 nA − 0.74 nA depending on the effective area to remove. A fine polishing procedure of the resulting cross sections was carried out on the sections, with a voltage of 30 kV and lower currents in the range 0.74 nA - 80 pA so that re-deposition phenomena in the cross section are very limited.

#### Image analysis and 3D reconstruction

All images were pre-processed with ImageJ (National Institute of Health, USA, http://imagej.nih.gov/ij). The images of the sequential cross sections shown in Figure 2, were collected as a stack, analyzed and processed with an open source tool chain based on Python (Python Software Foundation, USA, http://www.python.org) scripts and tools. The image stack was cropped, filtered and down-sampled. The isotropic resolution in x, y and z amounts to 38.5 nm. The reconstructed data are visualized with Blender. (Blender Foundation, Netherlands, http://www.blender.org).

